# Wrinkle-like structures emerge from matrix complementarity and mechanical discontinuity in heterogenous biofilms

**DOI:** 10.64898/2025.12.18.695073

**Authors:** Adrien Sarlet, Anja K. Ehrmann, Namiko Mitarai, Liselotte Jauffred

**Affiliations:** The Niels Bohr Institute, University of Copenhagen, Jagtvej 155 A, DK-2200 Copenhagen N, Denmark; DTU Bioengeenering, Technical University of Denmark, Søltofts Pl. Building 221, 2800 Kgs. Lyngby, Denmark

**Keywords:** Biofilm, Escherichia coli, morphology, multi-species, curli, phosphoethanolamine-cellulose, amyloid

## Abstract

Biofilms are dynamic communities of microorganisms encased in a self-produced extracellular matrix. These resilient structures pose challenges across nearly all human activities, from healthcare to industry. The mechanical behaviour of a biofilm is shaped by the heterogenous composition of its matrix, both spatially and chemically. In turn, these mechanics influence the biofilm’s architecture at micro- and macroscopic scales, driving its complexity and adaptability. Morphologically, this is reflected in mechanical deformations of the biofilm known as wrinkles. In nature, biofilms often host different species of bacteria, allowing for a great diversity of matrix components. Here, we use a combination of two *Escherichia coli* strains as a model for a multispecies biofilm in which each bacterial strain produces one of two complementary matrix fibres: an amyloid protein (curli) or a polysaccharide (phosphoethanolamine-cellulose). Using fluorescence microscopy, we confirm that the two bacterial strains rapidly segregate into isogenic sectors, decreasing local heterogeneity. Furthermore, we show how wrinkles form, both in the homogenous central region of the biofilm, as well as at the boundary between sectors (i.e. where the two matrix producers co-localize). Finally, we show that increasing strain intermixing via the addition of bacteriophages results in thicker, taller wrinkles, irrespective of whether the two fibres are produced by two different strains or co-produced by the same bacteria.

## INTRODUCTION

Biofilms form when microorganisms attach to a surface, divide, and embed themselves in a self-secreted extra-cellular matrix of polymeric substances (e.g. polysaccharides, proteins, lipids, and extracellular DNA). Growth in a biofilm structure collectively provides mechanical stability (1), enables nutrient and signal exchange (2, 3), protects bacteria against phage predation (4, 5), and protects against antibiotics (6, 7). Moreover, chemical treatments can either reinforce or weaken the mechanical stability of a biofilm depending on the matrix composition (8). Biofilms are assumed to account for the majority of prokaryotic biomass on Earth (9). They are involved in numerous diseases (10) and, more generally, negatively impact most sectors of human industries (11).

In *Escherichia coli* in particular, the extra-cellular matrix is mainly composed of curli amyloid fibres and phosphoethanolamine-cellulose (pEtN-cellulose) (12–14). The closely related *Salmonella spp*. is also known to produce both curli and pEtN-cellulose (14). Other Enterobacteriaceae were shown to produce curli and/or cellulose (15) while possessing the genes necessary for their pEtN-modification (16). All in all, the distribution of the genes responsible for the production of these two fibres is much more widespread than the few species in which their production was evidenced (16, 17). From *E. coli* populations producing only one of the two fibres, we know that curli-containing biofilms are stiffer (twice the Young’s modulus [Supplementary Note 1]) than those containing only pEtN-cellulose (18). More broadly, polysaccharides are known to drive biofilm colony expansion and mechanical properties (19). Together, these matrix fibres form a composite material that can withstand a higher compressive load than its individual components (18).

When surface attached communities grow, bacteria divide and push against each other. This leads to a build-up of mechanical tension in the community, which can be relieved by the formation of chiral swirls (20), new cell layers (21), bacterial motility (22), or by self-driven ordering of elongated cells (i.e. active nematics) (23, 24). Tension can also be released by the formation of *wrinkles*, which are generally understood as a mechanical buckling instability (25, 26). In other words, when compressive forces (arising from internal stresses) exceed a critical threshold, the biofilm undergoes out-of-plane deformation, resulting in the formation of wrinkles. Throughout this study, we use the term wrinkle to describe undulations in the surface of the biofilm, including those detaching from the surface (i.e. delamination) and with a length much larger than the width. Although environmental conditions leading to wrinkle formation differ between species, certain factors seem to be decisive, namely nutrient and oxygen availability (27), as well as agar concentration (proxy for substrate elasticity and friction). Specifically, as the outward expansion of the biofilm is opposed by bacteria-substrate friction, compressive stress is building up and the biofilm releases this stress by wrinkling (26, 28, 29). Therefore, in *E. coli*, increasing the concentration of agar generally promotes wrinkling (30). Although many parameters are known, a clear understanding of the cues that govern the onset and dynamics of wrinkling remain elusive, especially in multi-species biofilms.

In nature, one species can benefit from the matrix of another (31), e.g. *Vibrio cholerae* can protect *E. coli* against the predator bacterium *Bdellovibrio bacteriovorus* (32) or phage T7 (33). However, there are also examples of bacteria cooperating in matrix-formation. Namely, complementary mutant strains of *E. coli* and *Salmonella typhimurium*, each producing one of the two curli subunits (CsgA and CsgB) (34). Species-specific regulation of colony morphology has also been documented in *Shewanella algae* (35). Furthermore, a mutant *Bacillus subtilis* and *Pantoea agglomerans* were found to form wrinkled biofilms when well-mixed. In this example, *B. subtilis* provided only the peptidic component TasA, while *P. agglomerans* provided the exopolysaccharide (36).

During colony formation, stochastic fluctuations and cell-cell mechanics drive co-cultured strains to segregate into large isogenic sectors while competing for nutrients and space (37). This large-scale patterning depends on relative growth rates (38, 39) and inoculation densities (37, 38) and can be influenced by intra-species interactions (40) and phage predation (41, 42). But little is known about how this affects subsequent matrix formation and wrinkling.

In this study, we ask how the underlying spatial organization of the bacterial colony controls 3D biofilm architecture. We investigate how local cooperation between curli and pEtN-cellulose producing populations impacts wrinkle formation.Using gene recombination, we prepared a pair of (otherwise similar) *E. coli* strains, each producing one specific matrix fibre. We used this combination of strains as model biofilms, where different strains contribute complementary fibres to the extracellular matrix. Combining laser-scanning confocal microscopy and 3D-image analysis, we mapped the biofilms’ macroscopic architecture. We evaluated the role of genetic heterogeneity, and found that wrinkles and wrinkle-like structures are not restricted to regions where curli and pEtN-cellulose producing strains are well-mixed. Instead, they can also form on the boundaries between isogenic sectors. Lastly, we added the virulent phage T6 to the system to distort the spatial patterning of the colony and investigate subsequent changes in wrinkling. Based on these results, we propose a mechanical model of wrinkle-formation in heterogenous biofilms.

## METHODS

### Media

#### M63+xyl

The 5× M63 salt solution was composed of 15 g/l anhydrous KH_2_PO_4_ (≥98.0%, P9791, Sigma-Aldrich), 35 g/l anhydrous K_2_HPO_4_ (≥99.0%, 60353, Sigma-Aldrich), 10 g/l (NH_4_)_2_SO_4_ (≥99.0%, 09978, Sigma-Aldrich), 2.5 ml 20 mM FeSO_4_ (≥99.5%, 44970, Sigma-Aldrich), 20 mM Na-Citrate (≥99.5%, 71402, Sigma-Aldrich). The M63 minimal agar was composed of 1× M63 salt, 1 µg/ml thiamine hydrochloride (≥99%, T4625, Sigma-Aldrich), 2 mM MgSO_4_ (≥99.5%, 63138, Sigma-Aldrich), 2 mg/ml xylose (≥99%, W360600, Sigma-Aldrich) and 15 g/l BD Bacto^™^ agar (10455513, Fisher Scientific) dissolved in Millipore water.

#### M9 agar

The M9 agar was composed of 2 g/l glucose (22720, Serva), 1× M9 salts, 2 mM MgSO_4_ (0168, JT Baker), 100 µM CaCl_2_ (2461, Chemsolute), 1× trace elements and 0.5 µg/ml thiamine hydrochloride (T4625, Sigma-Aldrich) and 15 g/l BD Bacto^™^ agar (10455513, Fisher Scientific) dissolved in Millipore water. 5× M9 salts consist of 34 g/l Na_2_PO_4_, 15/g l KH_2_PO_4_, 2.5 g/l NaCl (27810.295, VWR) and 5 g/l NH_4_Cl (09711, Fluka). 2000× trace elements consist of 37 mM FeCl_3_, 7 mM ZnSO_4_, 2.3mM CuSO_4_, 5.9mM MnCl_2_, 2.5mM CoCl_2_, 1.6mM Na_2_EDTA, pH8. 50 µM NiCl_2_ was added when appropriate for counter-selection against *tetA*.

#### LB agar

The regular LB agar was composed of 10 g/l Gibco^™^ Bacto^™^ tryptone (16279751, Fisher Scientific), 5 g/l Gibco^™^ Bacto^™^ yeast extract (16279781, Fisher Scientific), 5 g/l NaCl (≥99%, S9888, Sigma-Aldrich) and 15 g/l BD Bacto^™^ agar (10455513, Fisher Scientific) dissolved in Millipore water. When specified, the LB agar was supplemented with 30 µg/ml kanamycin sulphate (≥95%, K1377, Sigma-Aldrich), 50 µg/ml spectinomycin dihydrochloride pentahydrate (≥62%, S-9007, Sigma) or 50 µg/ml tetracycline hydrochloride (T7660, Sigma-Aldrich).

#### Salt-free agar

The salt-free agar was composed of 10 g/l Gibco^™^ Bacto^™^ tryptone (16279751, Fisher Scientific), 5 g/l Gibco^™^ Bacto^™^ yeast extract (16279781, Fisher Scientific) and 15 g/l BD Bacto^™^ agar (10455513, Fisher Scientific) dissolved in Millipore water. When specified, the salt-free agar was supplemented with 30 mg/l Direct Red 23 (≥30%, 212490, Sigma-Aldrich) or 40 mg/l Congo Red (≥35%, C6767, Sigma-Aldrich), previously filter-sterilized and added to the melted agar immediately before pouring.

### Bacterial strains and culture

#### Background

MS613 (43) is a MG1655 K-12 strain producing curli amyloid fibres but not pEtN-cellulose (Cur^+^) due to an early stop codon in *bcsQ* in *E. coli* K-12 strains. AR3110 (12) is a W3110 (i.e. K-12) derivative in which *bcsQ* was repaired, thereby restoring pEtN-cellulose production.

#### AS011

From MS613, a xylose auxotrophic mutant was created by P1 transduction from the *xylA* mutant of the Keio collection (JW3537) (44), *xylA* being distant from *bcsQ* by approximately 35 kbp, followed by selection on kanamycin. The resulting xylose auxotroph was then *repaired* by P1 transduction from AR3110, followed by selection on M63 + xylose medium. The resulting strains were subsequently inoculated on salt-free agar for biofilm growth. One of those strains, AS011, showed similar wrinkling to that of AR3110. Thus, that strain’s *bcsQ* was restored simultaneously with *xylA* and therefore, it produces both curli fibres and pEtN-cellulose (Cur^+^pCe^+^). The P1 transduction protocol was adapted from (45, 46).

#### AS012

From AS011, *csgBA* was mutated by P1 transduction from RMV612 (20), which produces neither curli nor pEtN-cellulose (∅), thereby yielding AS012, which produces pEtN-cellulose but not curli (pCe^+^).

#### Introduction of fluorescent markers

For each of the four strains MS613, RMV612, AS011 and AS012 variants were constructed expressing either *gfp* or *rfp*. The fluorescent reporter genes were integrated into the genome of the strains following the protocols for λ-Red recombineering and *tetA* dual-selection protocol as described by Bayer et al. (47). Briefly, strains were first transformed with pSIM19 (48). The expression of the λ-Red proteins was induced by 20 min incubation at 42 °C. The cells were made electrocompetent by three rounds of washing in ice-cold water. 4 ng/µl gel-purified dsDNA or 400 nM ssDNA oligonucleotide were used for the transformation (final concentration in 50 µl competent cells). In the first step, a J23100-*gfp-tetA* dsDNA cassette, amplified from the genomic DNA of the strain SEGA007 (47) was integrated into the *glmS-pstS* intergenic region of all strains. Successful recombinants were selected on LB agar containing 50 µg/ml tetracycline. Candidate colonies were restreaked at least once on LB + tetracycline and subsequently verified by colony PCR and Sanger sequencing (Eurofinns Genomics, Germany). In order to exchange the *gfp* and *tetA* genes for *rfp*, or to remove *tetA* for a marker-less labelling with *gfp*, a second round of recombineering and *tetA* counter-selection was performed. Cells were transformed with either a *rfp* dsDNA fragment amplified from the genomic DNA of strain SEGA-007-*rfp* (47) or a ssDNA oligonucleotide containing the appropriate homology sequences to remove *tetA* from the J23100-*gfp-tetA* cassette. To select for the correct recombinants, cells were first recovered in LB medium overnight, washed in 1× M9 salts and a dilution row plated on M9 glucose plates supplemented with 50 µM NiCl_2_. Candidate colonies were restreaked twice on LB agar and verified by colony PCR and Sanger sequencing.

#### Biofilm culture

A liquid culture was inoculated from a single colony and grown overnight in LB medium. The optical density of the resulting bacterial suspension was measured at 600 nm (OD_600_). The bacteria were diluted to reach a concentration corresponding to OD_600_ = 0.4. Four drops of 2 µl each were typically inoculated onto a salt-free agar Petri dish. The resulting biofilms were sealed with Parafilm ^®^ and grown at 28 °C for 5 days. For biofilms inoculated from a mix of two different bacterial strains, each liquid culture was brought to OD_600_ = 0.4 before mixing them in the indicated proportion. Immediately before imaging, one biofilm was cut out from the agar and placed on a glass slide. The glass slide was flipped because the fluorescence and confocal microscopes were inverted.

#### Phage infection

The biofilms grown in the presence of T6 phages were inoculated as described above. After 6 h of incubation at room temperature, 1 µl of T6 phage suspension (6 · 10^8^ pfu/ml) was spotted either 1 mm away from the dried bacterial drop (side inoculation), or was added on top of the dried bacterial drop (centre inoculation). The method was derived from (41, 42). The biofilms were then incubated at 21 °C (room temperature) for 10 days before imaging. The phage suspension was diluted in salt free LB broth from a phage stock solution prior to infection.

Note that the phage experiments were performed under different growth conditions than the other experiments, following the protocol by Ruan et al. closely (41). They were also performed in a different lab facility using different stocks to prepare the growth medium.

To test susceptibility of strains to phage T6, liquid cultures of all bacterial strains were inoculated in LB medium from overnight cultures. Cultures were incubated at 37 °C until they reached exponential growth. 100 µl of bacterial culture were then mixed with 3 ml top agar (8 g/l NaCl (27810.295, VWR), 6 g/l BD Bacto^™^ agar (10455513, Fisher Scientific)) and poured on top of a fresh LB agar plate. After the top agar solidified, 2 µl of concentrated phage stock solution were added as a drop on top of the agar. Plates were incubated at 37 °C.

The possibility of phage T6 infection in salt free LB medium was evaluated by measuring OD_600_ growth curves of RMV612 (∅) in a FLUOstar Omega microtiter plate reader (BMG LABTECH, Ortenberg, Germany). After an overnight preculture in standard LB broth, RMV612 was diluted 1:250 into fresh salt-free LB broth (without agar) and incubated at 37 °C under shaking. When reaching exponential phase (OD_600_ ≈ 0.3 − 0.4) the bacteria were diluted again 1:10 into a 96-well microtiter plate containing salt-free LB broth and incubated at 37 °C for 45 min. Then, dilutions of a phage stock prepared in salt-free medium were added, the plate sealed with a breathable film and OD_600_ was measured every 5 min over the course of 20 h. The control samples were grown in LB medium supplemented with 10 mM MgSO_4_. The equivalent amount of salt-free medium was added to the phage-free control samples.

#### Cross-section

Following 5 or 10-day growth, 20 µg/ml chloramphenicol (≥98%, C0378, Sigma-Aldrich) was added to the Petri dish containing the biofilm to be sectioned. Prior to sectioning, the biofilm (and surrounding agar) was cut out and placed in a small Petri dish. There, the biofilm was embedded in agar also supplemented with 20 µg/ml chloramphenicol: ∼12.5 ml 50 °C agar was poured into the dish without directly pouring on the biofilm. The role of chloramphenicol is to prevent further colonisation of bacteria into the newly poured agar. Once the top agar had solidified, the biofilm was cross-sectioned with a scalpel, flipped, and transferred into a glass-bottomed dish for imaging. The cross-sectioning procedure was simplified from (30).

### Imaging and analyses

#### Brightfield imaging

Brightfield images with and without Congo Red, were acquired using a Reshape Imaging Device, Model 3.1a, Firmware 1.0.103 (Reshape Biotech, Copenhagen, Denmark) using the integrated 12 MP camera (Reshape Biotech) and top light illumination without optical magnification. Image analysis was performed via the Reshape Discovery Platform (2025).

#### Fluorescence microscopy

We used a Nikon ECLIPSE Ti microscope (Nikon, Tokyo, Japan) equipped with a 4× air immersion objective (Nikon, Plan Fluor, 4×/0.13, ∞/1.2 WD 16.5) and paired with an Andor Neo camera (Andor, Belfast, UK). GFP and RFP were excited by an Hg lamp using the FITC (FITC Filter Cube Set, Olympus, York, UK) and Texas Red cubes (Texas Red^™^ Filter Cube Set, Nikon, Tokyo, Japan), respectively. The tiles were stitched (20% overlap, pixel size: 1.625 µm) automatically upon acquisition by the microscope’s native software: NIS-Elements Advanced Research (4.13.03, Nikon).

#### Confocal scanning microscopy

We used a Leica DMI6000 CS SP5 (Leica, Wetzlar, Germany) with the Ar (488 nm) and HeNe (543 nm) laser lines sequentially to excite GFP and RFP, respectively. We used a 5× air objective (Leica, N Plan,5× /0.12 PH0) with a focal depth of ∼1.5 µm. We used *z*-steps of 0.5 µm, pinhole diameter of 88.44 µm, and scan speed of 400 Hz with line averaging of 3 (no frame averaging). For the cross sections, we used a 40× (Leica, N Plan, 40× /0.55) air objective with a focal depth of ∼0.5 nm and a pinhole diameter of 154.41 µm, but same scan speed, *z*-steps, and line averaging (no frame averaging).

#### Height maps and profiles

Using custom-made Matlab (Matlab R2023, The MathWorks, Inc) procedures and starting from *z*-stack images with two channels (GFP and RFP), the maximum intensity projection over the (*z*)-dimension of each channel was computed individually. Both projections were normalised so that their intensities could be compared. Every pixel of the image was assigned one colour (green or red) by comparing the two projections for that pixel. Then for every pixel in (*x, y*) in the original *z*-stack, the *z*-coordinate of the maximum intensity pixel was determined in the colour channel previously assigned to that pixel. This procedure relies on the maximal fluorescence intensity to originate from air-cell interfaces and subsurface attenuation to be limited due to optical confinement in dense biofilms. However, this method could give rise to a systematic subsurface bias and should, thus, be limited to relative height measurements (49). Then all *z*-coordinates were translated to the minimum value of the pixel and converted to metric units. This resulted in an (*x, y*)-image mapping the height of the brightest pixels: *h*-map. For wrinkle profiles, the *h*-map was used to find maximum wrinkle heights in pixel *i* over the length of the wrinkle: 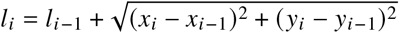.

#### Distances from wrinkle ridges to closest sector boundaries

The wrinkle ridge structures were extracted from *h*-maps using Gaussian smoothing (smoothing parameter of 4), Hessian-based ridge detection (threshold parameter 0.08), and skeletonization, where small objects below 100 pixels were removed. In parallel, sector boundaries were extracted from binarized maximum intensity projection over the (*z*)-dimension of the GFP-channel by removing small objects (less than 75-330 pixels), applying morphologic closing with a disk (radius of 1-3 pixels), and using the Canny edge detection filter. For each ridge coordinate, the Euclidean distance to the nearest sector boundary point was computed by using a k-d tree nearest-neighbour search with *k* = 1, and the histogram of the distance was created for each image, using the conversion of 6.0567 µm per pixel.

#### Strain occupancy

The images recorded from RFP-channels had generally lower intensities than the GFP channel. Therefore, when optimizing for full colony views, we underestimated the RFP in the thin bacterial periphery, compared to GFP [Figure 4]. To verify that this did not reflect a real attenuation, we did zoom-ins on the periphery [Supplementary Figure S6].

## RESULTS

### pEtN-cellulose reconstituted in curli-producing *E. coli*

Starting from a standard curli-producing but cellulose deficient (Cur^+^) *E. coli* K-12 strain, MG1655, we repaired the production of pEtN-cellulose by P1 transducing *bcsQ* from AR3110, in which it had been previously repaired (12). The resulting strain produces both curli and pEtN-cellulose (Cur^+^pCe^+^). From there, we mutated the curli producing genes *csgBA* by P1 transduction from the strain producing none of the two fibres (∅) (20), to create a pEtN-cellulose-only strain (pCe^+^) [Figure 1A and Table 1].

**Table 1:** Strains used in this work.

| Strain | Name | Genotype | Matrix composition | References |
| --- | --- | --- | --- | --- |
| MS613 | Cur<sup>+</sup> | MG1655 K-12 (F- λ- *ilvG*- *rfb*-50 *rph*-1) | curli | (22, 43) |
| RMV612 | ∅ | MG1655 *csgAB::kan* | / | (20) |
| AS011 | Cur<sup>+</sup>pCe<sup>+</sup> | MS613 *bcsQ*+ | curli, pEtN-cellulose | This work |
| AS012 | pCe<sup>+</sup> | AS011 *csgAB::kan* | pEtN-cellulose | This work |

**Figure 1:**
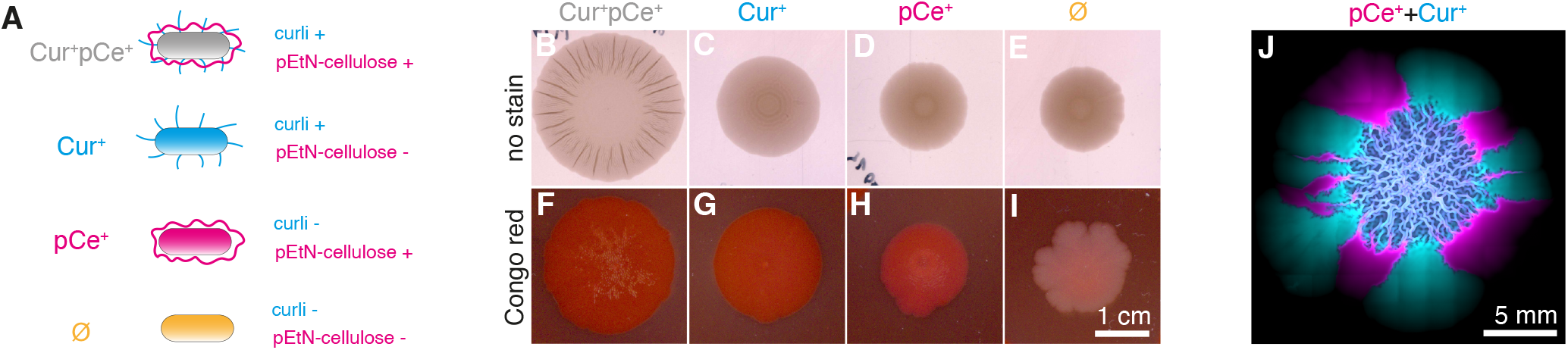
pEtN-cellulose reconstituted in curli-producing *E. coli*. *E. coli* biofilms grown for 5 days on salt-free agar (B-E) and on salt-free agar supplemented with Congo red (F-I). **A**: Illustration of the strain set used in this study. **B+F:** AS011 producing both curli and pEtN-cellulose (Cur^+^pCe^+^); **C+G:** MS613 producing only curli (Cur^+^); **D+H:** AS012 producing only pEtN-cellulose (pCe^+^) and **E+I:** RMV612 producing neither curli, nor pEtN-cellulose (∅). Scale bar: 1 cm. Note in particular the difference of shade between H and I, which otherwise appear identical in D and E. **J:** Pseudo-coloured fluorescence image of a two-strain biofilm (pCe^+^+Cur^+^) initially mixed in a 1:1 ratio grown for 5 days on salt-free agar. Scale bar: 5 mm.

Grown on salt-free agar, Cur^+^pCe^+^ biofilms exhibit long radial wrinkles [Figure 1B] and a larger diameter than biofilms formed by other strains [Supplementary Figure S1]. Cur^+^ biofilms form a previously described (12) characteristic ring pattern around the central region [Figure 1C, Supplementary Figure S2] but both the pCe^+^ strain and the ∅ mutant, producing none of the fibres, have smooth phenotypes [Figure 1D, E]. We confirmed the production of pEtN-cellulose in the pCe^+^ strain by supplementing the salt-free agar with Congo red, which binds both amyloid fibres and cellulose (12, 50) [Figure 1H, I]. Interestingly, the presence of Congo red also influences the biofilm phenotypes: The Cur^+^pCe^+^ biofilm showed shallower wrinkling [Figure 1F], while the pCe^+^ biofilm exhibited dense wrinkles in the central region, otherwise absent without Congo red [Figure 1H]. A weakening of the interaction between curli and pEtN-cellulose could explain the relatively smoother phenotype in the Cur^+^pCe^+^ strain. Such an effect by Congo red was already reported in *E. coli* UTI89 by Reichhardt *et al*. (50), who also reported increased wrinkling in their pEtN-cellulose-only strain in the presence of Congo red, due to a higher affinity of the dye for pEtN-cellulose than for curli.

We inoculated a suspension of the Cur^+^ and pCe^+^ strains (each labelled with a unique fluorescent reporter) in equal proportions. Once the colony expanded beyond the well-mixed area of the inoculation droplet, the population segregated into large isogenic sectors (38), each formed by a strain with the ability to produce one of the matrix fibres [Figure 1J]. As segregation is the result of small variations at the fast-growing colony front, this happens before the onset of matrix production. Specifically, the fibre production starts 48-72 hours after entry into stationary phase (5, 33). Therefore, we can use this system to test how a given spatial patterning affects the overall biofilm morphology. A zoom-in on the central region shows dense wrinkling and a pattern of curli and pEtN-cellulose producers (cyan and magenta) mixed at subwrinkle scales [Supplementary Figure S3A]. The corresponding height map (*h*-map) emphasises the surface structure of the complex architecture in the mixed biofilms [Supplementary Figure S3B-C]. We verified that curli and pEtN-cellulose fibres were present in both mixed and mono-strain biofilms by imaging biofilm cross-sections stained with Direct Red 23 [Supplementary Figure S4]. We found a strong stratification as characterized in a very similar system (51) and confirmed the presence of pEtN-cellulose within the wrinkle cavities and on the ridges. Based on this confirmation and to support readability, we will henceforth refer to strains capable of producing these fibres as *producers*. However, while remembering that matrix production varies in different layers of the biofilm and that certain subpopulations might not produce any fibres at all.

### Wrinkle-like structures are formed on the boundaries of curli and pEtN-cellulose-rich regions

Outside of the homogenous central region, the two strains are only in contact at the boundary between their respective isogenic sectors [Figure 2A]. We verified that there was no strain stratification by cross-sectioning the biofilm [Figure 2B]. Despite this strong demixing of genotypes, the biofilm formed radial wrinkle-like structures at the sector boundaries exclusively. We did not observe any wrinkles in isogenic sectors, which indicates a synergistic interplay between the strains, each producing one matrix fibre. Morphologically, two types of radial wrinkle-like structure were observed. First, wrinkles formed on the boundaries between large adjacent sectors of mono-strains, each producing one of the fibres [Figure 2C]. Alternatively, wrinkles formed around a thin pCe^+^ sector enclosed between two Cur^+^ sectors [Figure 2D]. We cross-sectioned sector boundaries of mixed pCe^+^+Cur^+^ biofilms (i.e. cross-sections perpendicular to wrinkles) to characterize the two different types of wrinkle-like structures [Figure 2E-F]. In both cases, the pCe^+^ bacteria (magenta) dominate the ridge (white arrows) of the wrinkles. We also found that at the sector boundary, the mixed pCe^+^ and Cur^+^ layer was present at the surface, while the pCe^+^ bacteria extended underneath [Figure 2E]. In both cases pCe^+^ filled up the cavity of the wrinkle [Figure 2E-F]. Staining with Direct Red 23 confirms the presence of both matrix fibres in the wrinkle-like structure [Supplementary Figure 4D]. The patterning of the two fibres in the wrinkle corresponds to the patterning of the two strains at the interaction interface, confirming the non-diffusive nature of the matrix fibres and the presence of pEtN-cellulose inside the wrinkle cavities.

**Figure 2:**
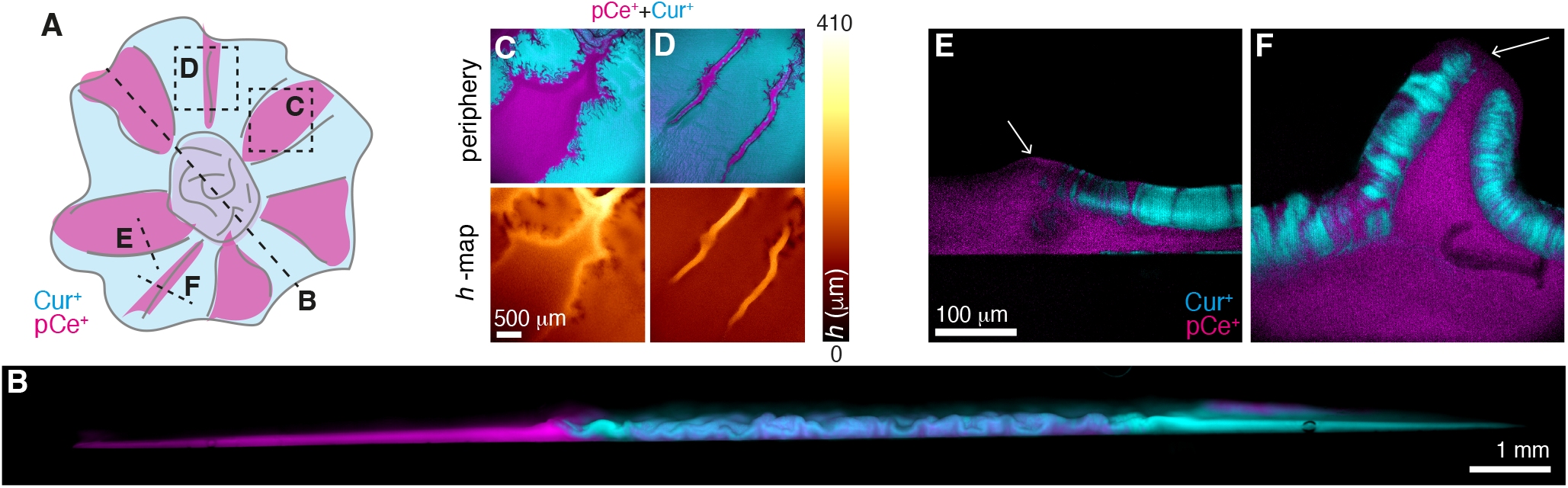
Wrinkle-like structures are formed on the boundaries of curli and pEtN-cellulose-rich regions. **A:** Sketch of a two-strain biofilm from a 1:1 mixed suspension (Cur^+^+pCe^+^). The letters indicate from where in the colonies the images in (C-D, black boxes) and the cross-sections in (B,E-F, black lines) were taken. **B:** Pseudo-coloured wide-field fluorescence microscopy of a cross-section through a biofilm grown from a 1:1 suspension of curli- and pEtN-cellulose-producing strains (Cur^+^+pCe^+^). Scale bar: 1 mm. **C-D:** Pseudo-coloured maximum-intensity projections (periphery) and the respective height map (*h*-map) from the outer region of the two-strain colony (Cur^+^+pCe^+^). Scale bar: 500 µm. **E-F**: Pseudo-coloured confocal imaging of cross-sections of wrinkle-like structures in the outer region, at the boundary between two larger sectors (E) and through a thin pEtN-cellulose sector (F). Scale bar: 100 µm.

### Wrinkle-like structures are formed in curli-sparse regions

As curli and pEtN-cellulose fibres possess distinct mechanical properties (18, 52), they contribute differently to *E. coli* biofilm architecture. Curli is considered to be the stiff – but brittle – component that is able to withstand a large compressive load. In contrast, pEtN-cellulose acts as a glue-like tension-bearing matrix component (53). We therefore investigated whether the wrinkle formation we observed on the boundaries between curli and pEtN-cellulose-rich regions requires both fibres or rather arises from the interruption of curli sectors. To do so, we mixed the Cur^+^ with the ∅ strain, which produces neither curli nor pEtN-cellulose. We found stronger wrinkling in the centre region of the pCe^+^+Cur^+^ biofilm than in the ∅+Cur^+^ biofilm [Figure 3A-C]. This indicates that ∅-dilution of the Cur^+^ strain creates a more viscous material capable of dissipating the stress. Nevertheless, when considering the segregated periphery of the biofilm, the thin sectors (pCe^+^ or ∅) squeezed between two Cur^+^ sectors rose in wrinkle-like structures in both cases [Figure 3D-E]. The structures had similar topography and maximal heights (*h* ≈ 100 µm), similar in height to the wrinkles of the pCe^+^+Cur^+^ mix’ central region [Figure 3A]. These results indicate that wrinkle formation at the sector boundaries of a multi-strain biofilm is the consequence of an interruption of rigid (curli-rich) sectors and does not necessarily require the presence of pEtN-cellulose. Therefore, wrinkling here is (at least partially) determined by the elastic properties of the biofilm.

**Figure 3:**
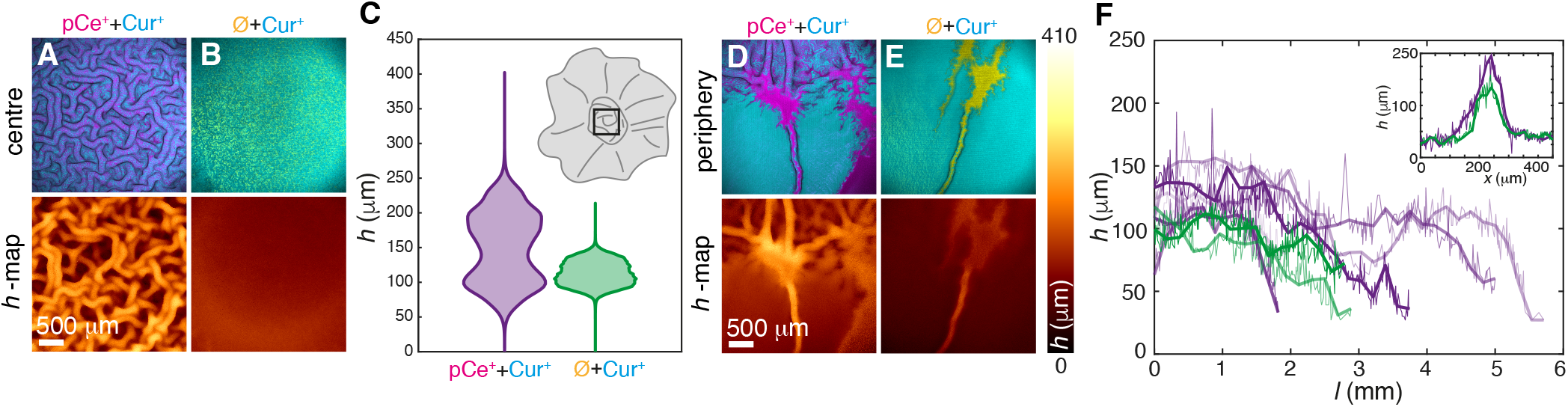
Wrinkle-like structures are formed in curli-sparse regions. **A-B:** Pseudo-coloured maximum-intensity projections (centre) and the corresponding height map (*h*-map) of a region-of-interest from the central region of two-strain colonies of (pCe^+^+Cur^+^) (A) and the curli-producer and the strain producing neither curli nor pEtN-cellulose (∅+Cur^+^) (B), see legends. Scale bar: 500 µm. Colour bar in (E) and note that the data in A is the same as in Supplementary Figure S3. **C:** Height distributions of the *h*-maps in (A-B). The inset is a sketch of the biofilms, where the central region-of-interest is indicated (black box). *N* = 262144 in both cases. **D-E:** Pseudo-coloured maximum-intensity projections (periphery) and the corresponding height map (*h*-map) of a region-of-interest from the outer region of the two-strain colonies (pCe^+^+Cur^+^) (D) and (∅+Cur^+^) (E), see legends. Scale bar: 500 µm. **F:** Maximum wrinkle heights, *h*, versus the length of the radial wrinkle, *l*,measured from the centre and outwards. Examples of wrinkles from two-strain colonies of pCe^+^+Cur^+^ (purple) and ∅+Cur^+^ (green), where the thicker full line is a spline-like smoothed curve-fit. Inset: *h* versus the axis perpendicular to the wrinkle, *x*, for pCe^+^+Cur^+^ (purple) and ∅+Cur^+^ (green). Colour scale corresponds to the main plot.

### Phage predation changes biofilm spatial organization through increased intermixing

We have established that biofilm morphology reflects the underlying multi-strain colony and that wrinkles form sub-sequent to growth-driven spatial organization. To quantify this, we calculated the distances from wrinkle ridges to sector boundaries in mixed biofilms (pCe^+^+Cur^+^) and found a strong correlation compared to the mono-strain Cur^+^pCe^+^ biofilm producing both fibres [Supplementary Figure S7-S8]. In the next step, we decided to manipulate this organization through bacteriophage infection. Indeed, Ruan and co-authors have previously demonstrated that predation by phage T6 on a growing colony slows down spatial segregation through increased cell-cell alignment (41). In other words, we used phage predation to enforce homogeneity into our otherwise heterogenous biofilm matrix. First, we confirmed that all strains were sensitive to phage T6 and that the phage could infect and spread in salt-free medium [Supplementary Figure S5]. Then we added phage T6 to a mixed (pCe^+^+Cur^+^) biofilm shortly after initial inoculation and observed how segregation was suppressed during range expansion [Figure 4A and Figure S7]. As a consequence, wrinkling ceased to be restricted to the central region and predefined sector boundaries. Instead, pCe^+^+Cur^+^ biofilm morphology resembled a Cur^+^pCe^+^ biofilm under phage T6 predation, characterized by large wrinkles with large separation (i.e. wavelength) [Figure 4B]. Specifically, the wrinkle expansion includes a broadening (in the surface plane) and a *h*-raise [Figure 4C] for the radial wrinkles, reaching approximately 500 µm in both directions [Figure 4D]. From cross-sections of these radial wrinkles, we confirmed that the composite material (i.e. pCe^+^ and Cur^+^ under phage T6 predation) retained diversity throughout the biofilm. This also resulted in different patterning of the wrinkles [Figure 4E, cf. Figure 2E-F]. Moreover, one large wrinkle often consisted of a small set of wrinkles [Figure 4E]. Probably resulting from individual wrinkles pushed toward each other by the adjacent flat regions (29). Alternatively, we also inoculated the phages a few mm away from the nascent colony. The result was that the phenotypical change (i.e. large wrinkles with large separation) was restricted to the area of contact with the phages [Figure 4F]. The above observations are consistent with phage T6 erasing faster-growing biofilm fronts, which prevents segregation of strains (41). However, it is worth noting that T6 predation did not shape our biofilm fronts to mono-layers of bacteria, in contrast to what was reported for similar *E. coli* strains (41). Instead, biofilms retained their volume [Supplementary Figure S6], indicating some protection (e.g. shielding) from matrix fibres.

**Figure 4:**
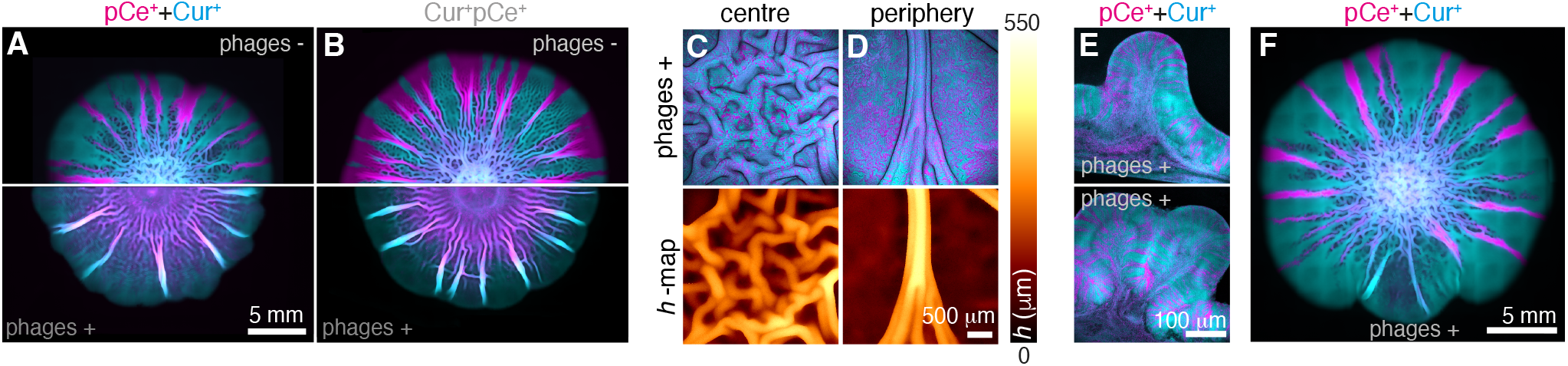
Phage predation changes biofilm spatial organization through increased intermixing. **A-B:** Pseudo-coloured fluorescence images of pCe^+^+Cur^+^ biofilm (A) and Cur^+^pCe^+^ biofilm grown of isogenic but coloured (magenta+cyan) strains (B). In both cases, they where grown for 10 days on salt-free agar with (phages+) or without phage T6 added (see legends). Scale bar: 5 mm. The change in phenotype of the negative controls (compared to Figure 1-4) is due to a change in growth conditions (see Methods). **C-D:** Pseudo-coloured maximum-intensity projections and their respective height maps (*h*-map) from the central region (B) and outer periphery (C) region of the two-strain colony (Cur^+^+pCe^+^) with T6 phages added (phages+). Scale bar: 500 µm. **E**: Pseudo-coloured confocal imaging of cross-sections of wrinkles in the outer region (periphery) in two-strain colonies (Cur^+^+pCe^+^) with T6 phages added (phages+). **F:** Pseudo-coloured fluorescence images of the two-strain biofilm (pCe^+^+Cur^+^) initially mixed in a 1:1 ratio grown for 10 days on salt-free agar with phage T6 added next to the biofilm (see legends). Scale bar: 5 mm.

## DISCUSSION

Biofilm-formation relies on the secretion of an extracellular matrix that constitutes a robust three-dimensional architecture. The resulting surface wrinkles depend on cell-cell interactions, substrate adhesion, and the mechanical properties of the matrix fibres. For example, curli and pEtN-cellulose have either synergistic or antagonistic effects on bacterial adhesion, depending on the substrate (54–56). Multiple studies have shown that the distribution of matrix component is heterogenous, even in isogenic systems (18, 57), and that heterogeneity arises from intricate regulation of matrix production (57). To capture this spatial heterogeneity, agent-based models have incorporated substrate friction, adhesion, and nutrient dynamics (58–60). In addition, small adhesion and nutrient variabilities were proposed to shape the global pattern (61).

Building on this framework, we sought to decipher heterogeneity arising from the spatial organization of multiple strains contributing to a biofilm, rather than from variations in gene expression. To this end, we investigated matrix complementarity by mixing *E. coli* curli- and pEtN-cellulose producers (Cur^+^+pCe^+^) with distinct mechanical properties (18, 53). The resulting surface-attached biofilms had a homogeneous composite centre and a heterogenous periphery composed of mono-strain sectors separated by well-defined boundaries.

In the homogeneous central region of the mixed biofilm, we observed intense wrinkling, even exceeding that of the strain producing both fibres (Cur^+^pCe^+^). While the initial deposition on the substrate causes an up-concentration of cells on the rim of the inoculation droplet (i.e., “coffee ring” (62, 63)), likely the inwards growth will enhances the compressive stress and the resulting wrinkling (29). However, wrinkling intensity was sensitive to subtle growth variations (compare Figure 1B and Figure 4A with Supplementary Figure S2). These variations may be reflected in substrate mechanics (64–66), signalling (3), cell death (25) or matrix production (57).

In contrast, in the heterogeneous peripheral region, wrinkle-like structures were restricted to boundaries between curli (Cur^+^) and pEtN-cellulose (pCe^+^) sectors, where complementary matrix fibres interact. Furthermore, under specific conditions, wrinkles formed on the boundary of sectors of the strain producing neither fibres (∅). These observations favours the conclusion that tension is building up in curli-rich regions of high rigidity and that this tension is released by spontaneous wrinkling in the more flexible (cellulose-rich) regions.

From the cross-sections, we found biofilms detaching from the substrate (i.e. delaminated), similar to what has been reported for *Pseudomonas aeruginosa* (67) and *Vibrio cholerae* (29). However, this matches the expectations for high substrate rigidity (1.5% agar) of delamination favoured over wrinkling (29). We also found pEtN-cellulose producers filled the wrinkle interior, making it unlikely that these wrinkles could serve as transport channels (12, 14, 68), as in *B. subtilis* (69). However, it remains unresolved whether cellulose-producing cells progressively squeeze between leaflet and substrate or occupy gaps created by curli-layer delamination. In either case, this suggests that there could be more pEtN-cellulose producers in the biofilm, than what appears from the top views [Supplementary Figure S3A].

Biofilms have been shown to protect against phage infection. Mature *E. coli* biofilms resist phage T7 via curli-mediated inhibition of phage transport and binding (5), and in *Staphylococcus epidermidis*, biofilm architecture strongly influences phage efficacy (70). Phages can also degrade extracellular fibres or exploit them as receptors (71–73). However, none of these activities have been observed for phage T6, and we did not test interactions between T6 and curli or pEtN-cellulose fibres as such (74). Instead, phage exposure was used to delay segregation and, thereby, affect wrinkling. Indeed, we found that mono-strain (Cur^+^pCe^+^) and mixed (pCe^+^+Cur^+^) biofilms had similar morphologies under infection. To our knowledge, this is the first demonstration of phage-induced changes in biofilm wrinkling. Following Ruan *et al*. (41, 42), we attribute this to changes in the directional order in the biofilm. While phage-mediated killing at the colony front increases alignment of rod-shaped bacteria (behind the front), diversity is retained and the merging of sectors is delayed. This indicates that the resulting composite matrix resembled that of mono-strain biofilms.

Directional elastic instability is a general feature of anisotropic thin films, where wrinkle orientation and spacing depend on the alignment of the material director relative to the compressive stress (75). In biofilms, orientational ordering of rod-shaped bacteria generates anisotropic stress distributions (76), correlating with aspect ratio (23), adhesion (24, 77), and matrix formation (in *V. cholerae* (76, 78)). Thus, phage-induced alignment likely creates mechanical anisotropy and may bias wrinkling toward fewer, more dominant radial modes. Phage infection may additionally alter stress-signalling pathways, controlling matrix production (3).

These observations were integrated within an extended mechanical framework, covering both the homogenous biofilm interior and the heterogenous periphery [Figure 5]. Mechanical *thin sheet* models (79) regard biofilms as flexible material expanding radially, while the adhesion of the film to its substrate generates friction in the direction of the biofilm centre (anti-parallel to expansion). This leads to a build-up of shear stress in the biofilm, relieved by wrinkling. This description covers the homogeneous biofilm interior [Figure 5A]. To predict wrinkling size and periodicity, homogenous biofilms have been described as continuous thin elastic plates and wrinkles as Föppl-von Kármán equations (based on Young’s modulus, Gauchy tensors, and Poisson ratio) (80, 81). Furthermore, adding additional thin plates with different elastic properties can account for colony-substrate friction, gradients (e.g. of air, humidity, and nutrients) (28, 29, 82), and viscous relaxation (83). To encompass the peripheral region, we propose to incorporate spatial heterogeneity as thin (curli-rich) plates interrupted by softer tension-bearing (pEtN-cellulose) regions [Figure 5B]. Within this heterogeneous elastic landscape, stress accumulates in stiff domains and is released at compliant interfaces, explaining the emergence of wrinkles at sector boundaries.

**Figure 5:**
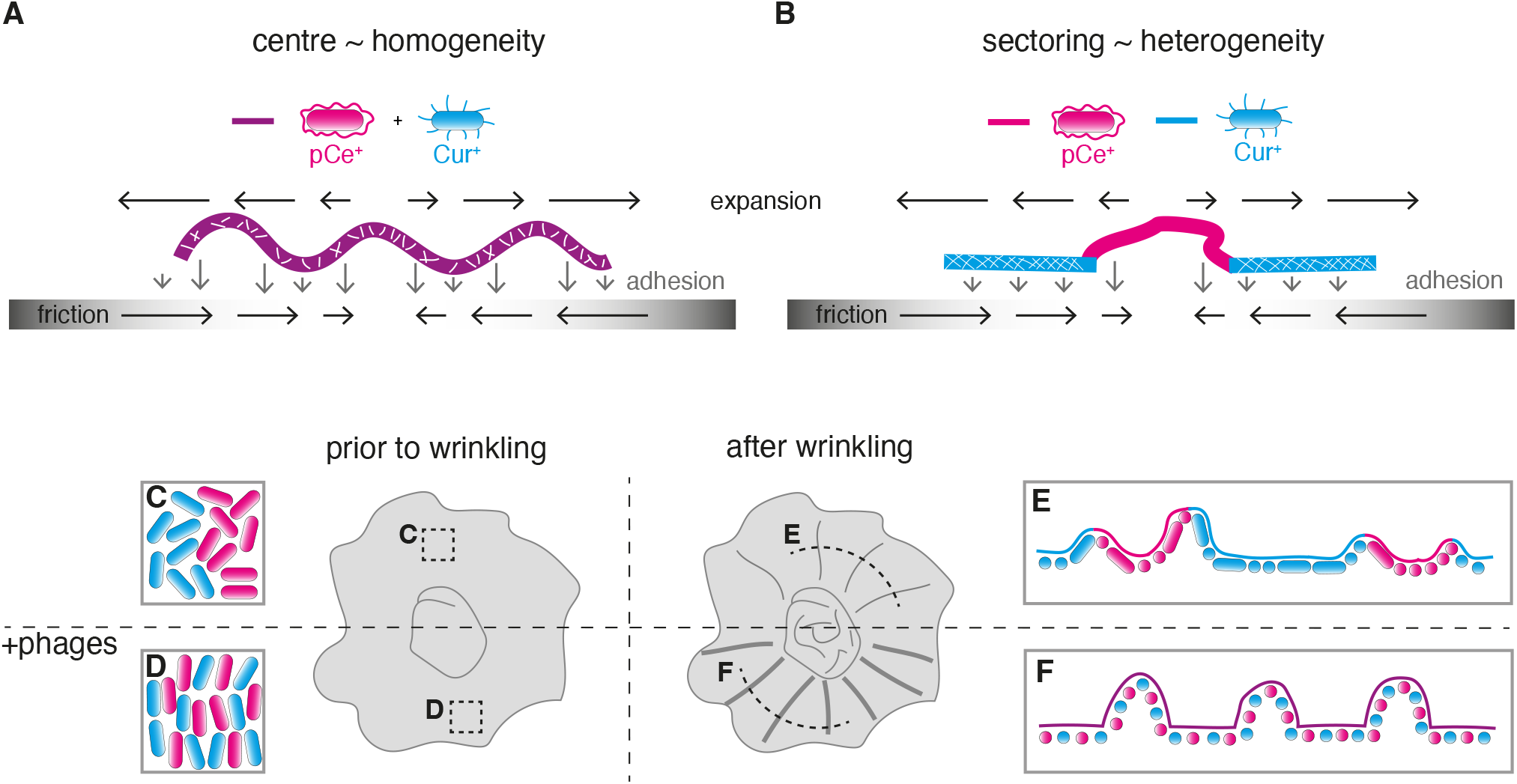
Wrinkling in homogenous and heterogenous biofilms. **A-B:** Illustration of wrinkle-formation in the homogenous central region (A) and on the heterogenous sector boundaries (B). The expansion of the biofilm is counteracted by the friction at the biofilm-agar interface (black arrows) and adhesion (gray arrows). The composite material of both strains and fibres (purple) is semi-rigid (dilute network of white lines) and the mono strain sectors of pEtN-cellulose (pCe^+^) and curli (Cur^+^) producers are soft or rigid (tight network of white lines), respectively. **C-F:** Conceptual model of orientational order prior to wrinkling (C-D) or after wrinkling (E-F) with phages (C+E) or without (D+F). The letters indicate from where in the colonies the top views (C-D, black boxes) and the cross-sections (E-F, black lines) were taken. Note that while the illustrations of the orientational order prior to wrinkling builds on earlier reports (41), the orientational order after wrinkling is an interpretation of the results of this study.

Phage-induced alignment further modifies this mechanical landscape by imposing directional elasticity and enhancing compositional homogeneity, thereby, shifting wrinkle periodicity and regularity [Figure 5C-F]. In this way, we provide a unified mechanical interpretation linking matrix complementarity, spatial organization, and phage-mediated re-organization to global biofilm architecture [Figure 5].

Our study establishes *Escherichia coli* as a versatile model for investigating wrinkle-formation in Gram-negative biofilms. By reconstituting the two essential matrix components (curli amyloid fibres and phosphoethanolamine-cellulose) we created a fully controllable two-strain system that mimics multi-species heterogeneity. This approach enables systematic explorations of how matrix composition and spatial organization govern mechanical instabilities such as wrinkling manifest in the macroscopic biofilm morphology. Given the growing clinical threat of Gram-negative biofilms (84), this model system provides a valuable platform to decipher the mechanics of biofilms and develop strategies to mitigate biofilm-associated problems.

## Acknowledgments

The authors thank Sine Lo Svenningsen and Bertil Kent Gummesson for access to the Keio collection and to their microbiology facilities, Cécile Bidan and Jeannette Steffen for providing the AR3110 strain, and Stanley Brown for his mentoring regarding the P1 transduction.

## Author Contributions

AS conceived the original idea. AS and AKE performed the experiments: both contributed to the strain engineering and to the imaging. AS did the data analysis. LJ and NM supervised the project. All authors discussed the results and contributed to the final manuscript.

