## Supplementary Information for "Wrinkle-like structures emerge from matrix complementarity and mechanical discontinuity in heterogenous biofilms"

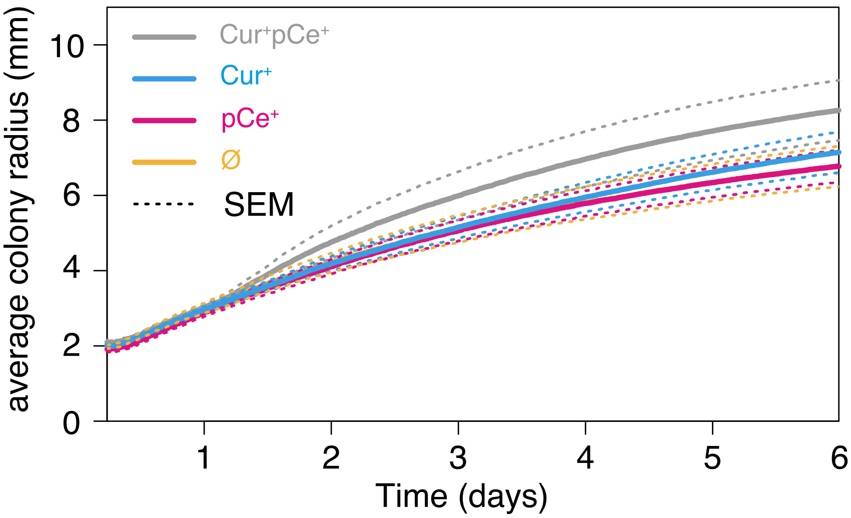


**Figure S1**. **Expansion rates of mono-strain colonies.**
The average colony radius versus time for the wild-type and all mutants individually. $N = 4$ an the dotted lines indicates ±SEM.

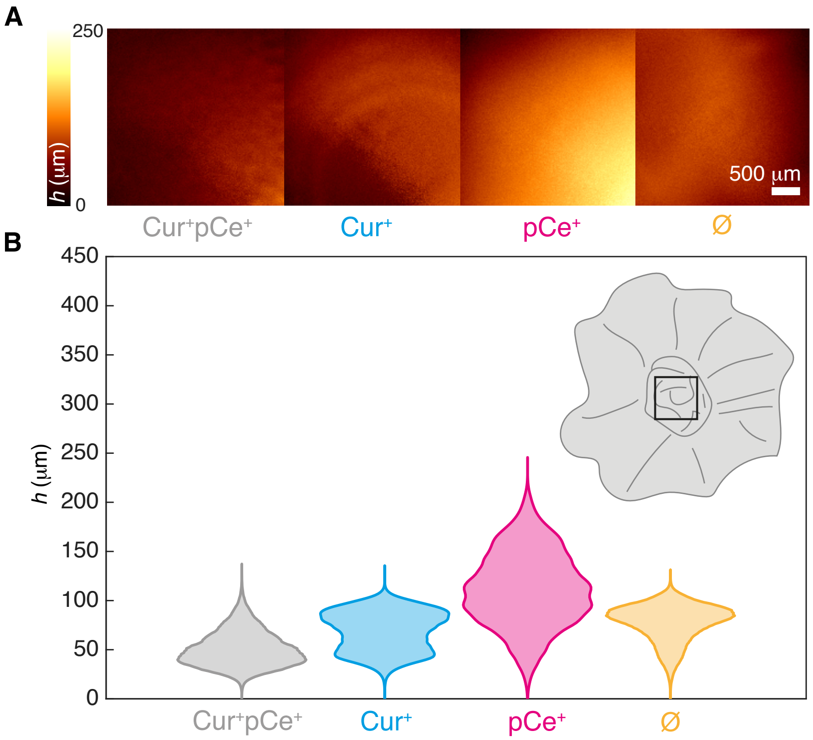


**Figure S2**. **Height maps of mono-strain colonies.**
**A:** Height maps of *E. coli* mono-strain biofilms grown for 5 days on salt-free agar (see legends). The scale bar:500 µm. **B:** Height distributions (violin plots) of the maps in (A). The inset is a sketch of the *E. coli* biofilm, where the central region-of-interest of (A) is indicated (black box). $N$ = 262144 in all cases and the experiment was repeated independently 1 time.

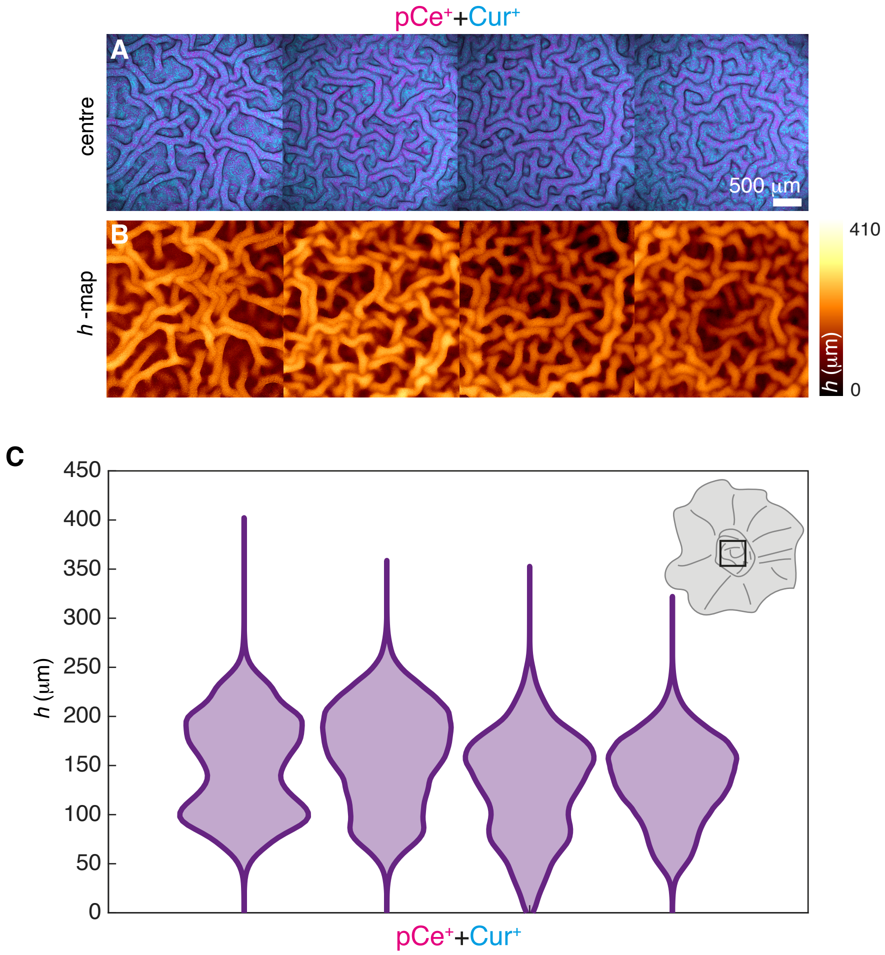


**Figure S3**. **Height maps of two-strain colonies.**
**A:** Pseudo-coloured maximum-intensity projections of region-of-interests from the central region (see inset of (C)) independent replicates of *E. coli* two-strain colonies ($Cur^{+}$+$pCe^{+}$). Scale bar: 500 µm. **B:** The according height maps (*h*-map). **C:** Height distributions (violin plots) of the maps in (A). The inset is a sketch of the biofilm, where the central region-of-interest of (A) is indicated (black box). $N$ = 262144 in all cases and the experiment was repeated independently 4 times.

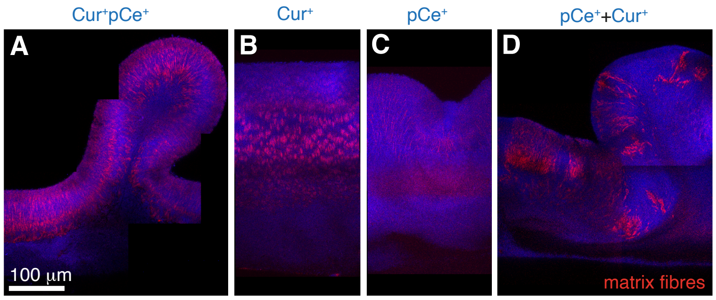


**Figure S4**. **Verification of matrix production**
**A-D:** Pseudo-coloured confocal imaging of cross-sections of wrinkles in the outer region of colonies from $Cur^{+}pCe^{+}$ (A), $Cur^{+}$ (B), $pCe^{+}$ (C), and the combination ($pCe^{+}$+$Cur^{+}$). All bacteria are GFP-producing (blue) and matrix fibres are stained with Direct Red 23 (red). Scale bar: 100 µm.

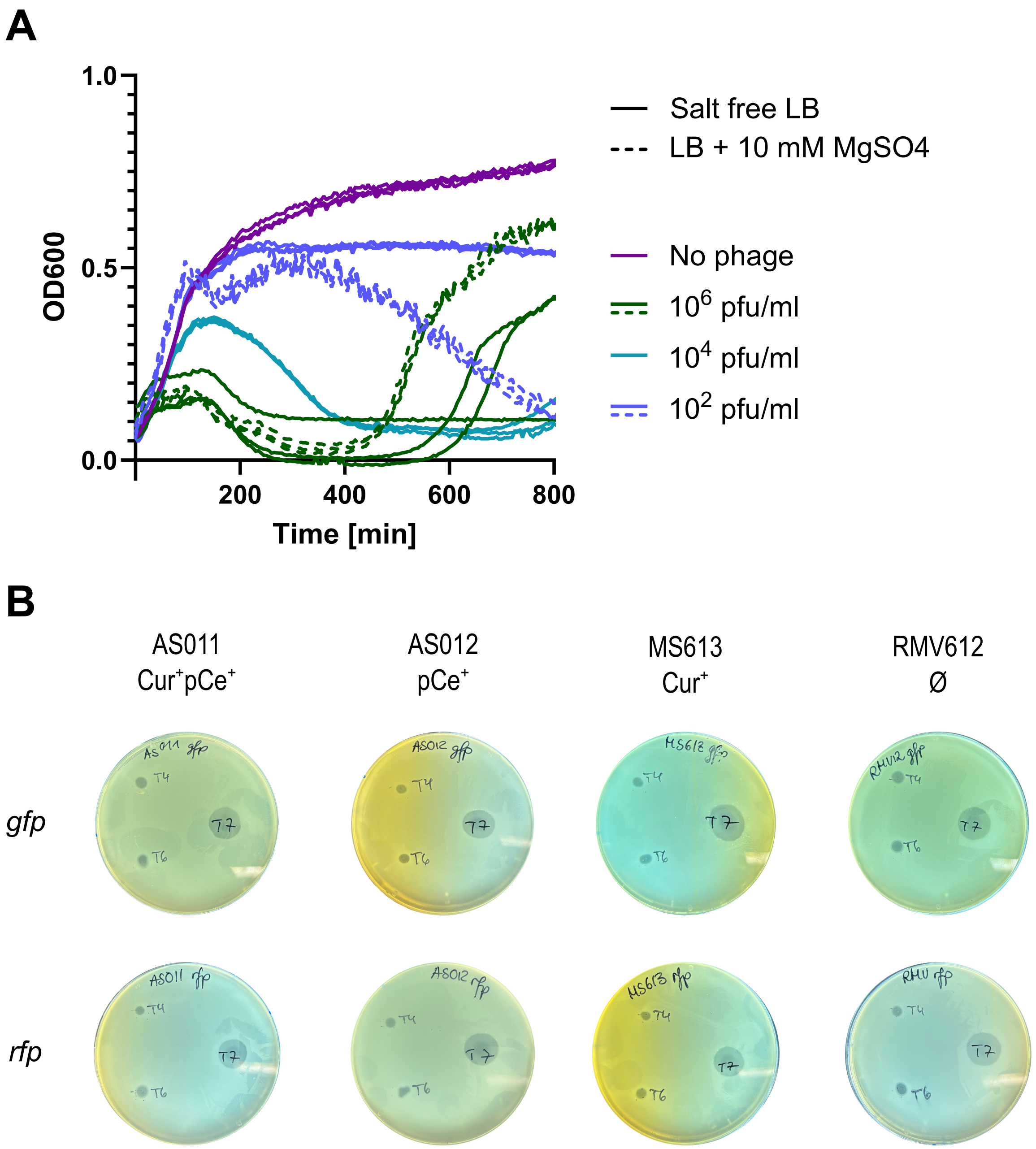


**Figure S5**. **Verification of phage T6 sensitivity and success of T6 infection in salt-free medium**
**A:** $OD_{600}$ growth curves of RMV612 (∅) in salt-free broth under infection by phage T6. Solid lines show samples grown in salt-free medium, dashed lines indicate control experiments in regular LB medium supplemented with 10 mM $MgSO_4$. The given phage titer refers to the final phage concentration in the samples at the start of the experiment. At low phage concentrations, infection is less efficient in the salt-free medium, but there is little difference between the two conditions at a sufficiently high phage titer. For all conditions three replicates are shown. Measurements were taken every 5 min. **B:** All strains used in this study are sensitive to phage T6 (as well as phage T4 and T7). Bacterial lawns were prepared by the agar overlay method and drops of concentrated phage stock were added on top of the solidified agar.

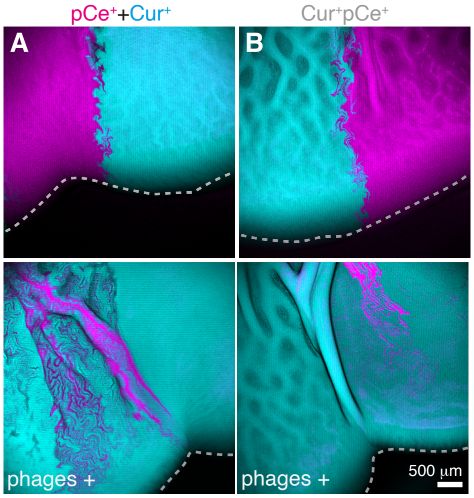


**Figure S6**. **Phage predation promotes mixing of matrix-producers but does not leave a mono-layered biofilm front.** Pseudo-coloured maximum-intensity projections of region-of-interests of colony fronts at the boundaries between two sectors. Punctuated lines (gray) correspond to fronts of expanding colonies. **A:** Mixed biofilm of curli ($Cur^+$) and cellulose-producing ($pCe^+$) without (top) or with phage T6 (bottom, side inoculation). **B:** Biofilm of the strain producing both fibres ($Cur^+pCe^+$) both with two different colours without (top) or with T6 phage (bottom). This corresponds to zoom-ins on biofilms mixed in a 1:1 ratio grown for 10 days on salt-free agar with phage T6 added next to the biofilm like in Figure 4F in the main text. Scale bar: 500 µm.

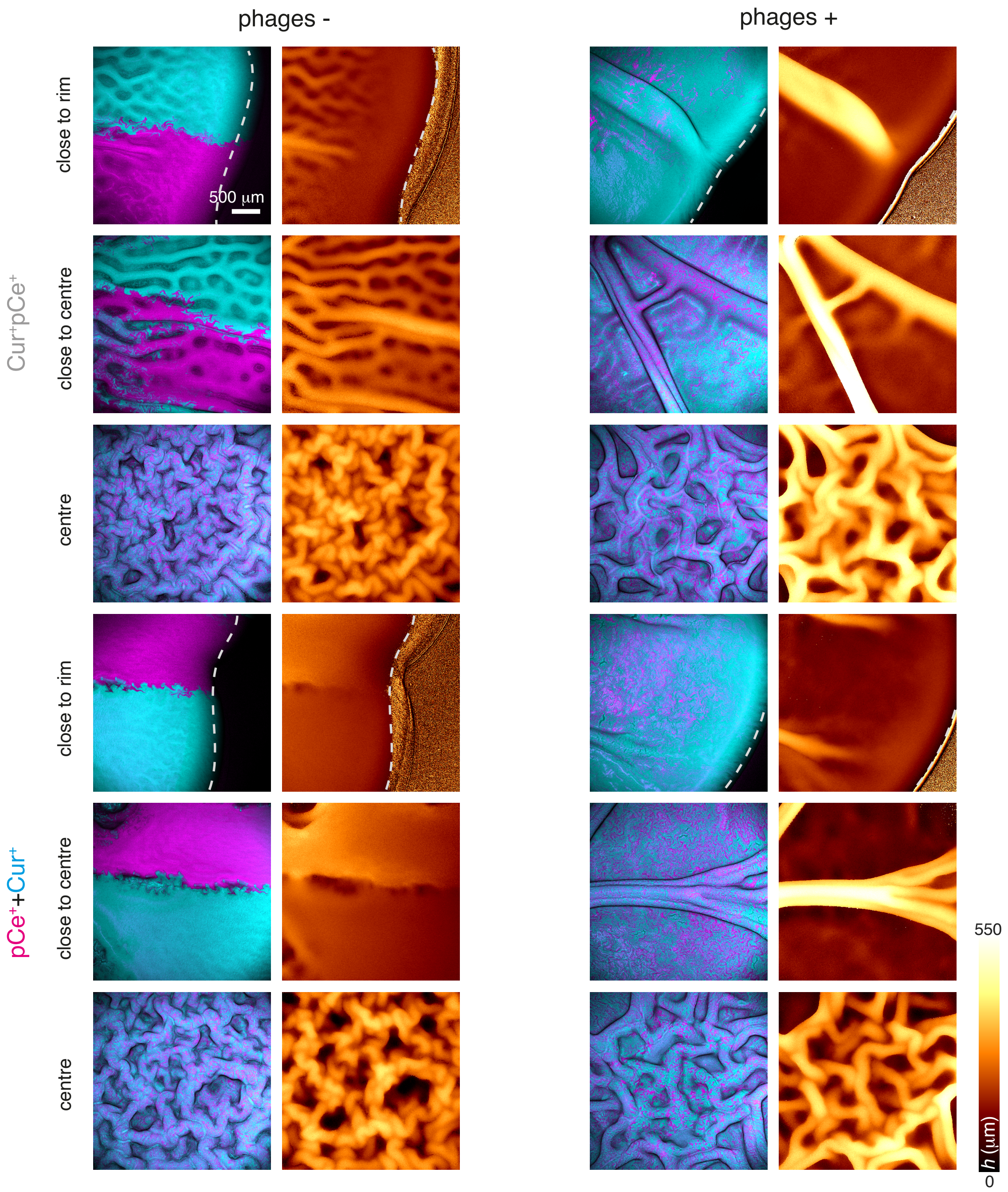


**Figure S7**. **Phage predation promotes mixing of matrix-producers.**
Pseudo-coloured maximum-intensity projections of region-of-interests from biofilm of the strain producing both fibres ($Cur^+pCe^+$) with two different colours and mixed biofilm of curli ($Cur^+$) and cellulose-producing ($pCe^+$) without (phage−) or with phage T6 (phage+). Note that some images are duplicates of images in Figure 5. Punctuated lines (gray) correspond to fronts of expanding colonies. Scale bar: 500 µm.

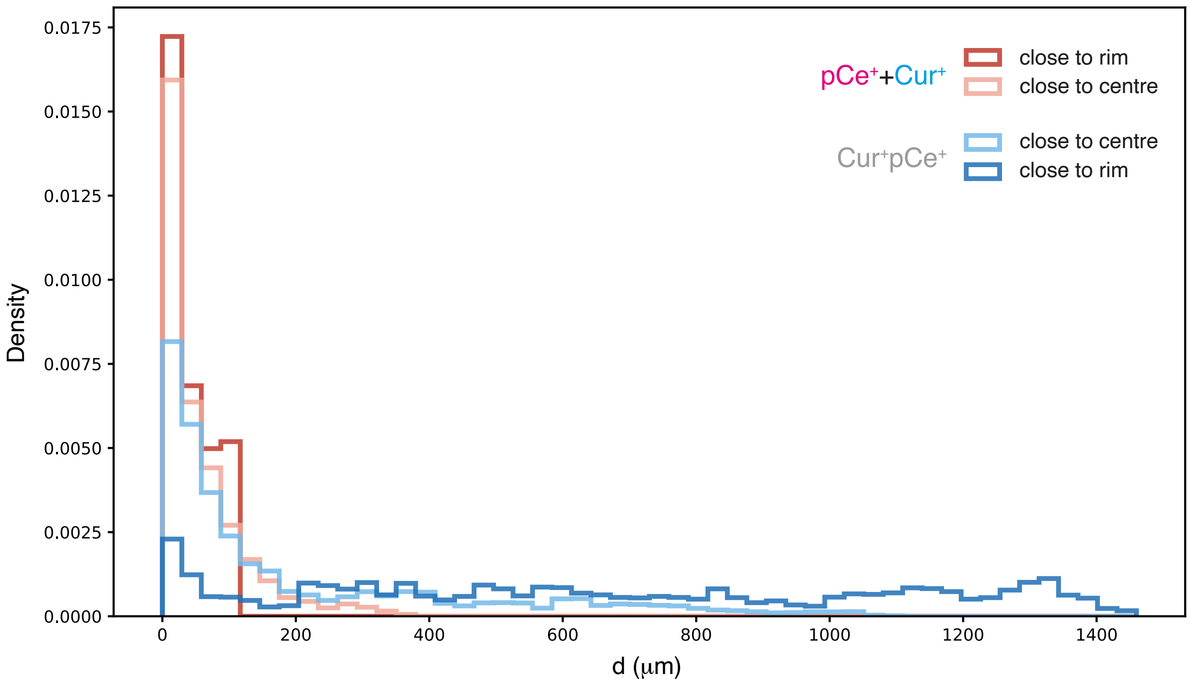


**Figure S8**. **Distances from wrinkle ridges to nearest sector boundaries.**
Density distributions of distances, *d*, calculated from the region-of-interests from biofilm of the strain producing both fibres ($Cur^{+}pCe^{+}$) with two different colours and mixed biofilm of curli ($Cur^{+}$) and cellulose-producing ($pCe^{+}$). Note that these values are calculated from the images in Figure S7, compare legends.

### SUPPLEMENTARY NOTE 1

From a very similar *E. coli* system (1), we got the median reduced elasticity, $E_r$, for both strains producing only one of the two fibres in question: 45 kPa and 23 kPa for $Cur^+$ and $pCe^+$, respectively. $E_r$ relates to Young's modulus, $Y$, as (2):

$$E_r \approx Y/(1-\nu^2), \quad (1)$$

where $\nu$ is the Poisson ratio in the range of 0.4-0.5 for typical biofilms, close to the incompressible limit ($\nu = 0.5$) (3, 4). Therefore, if we assume negligible differences between $\nu$ in the mono-strain biofilms, we get the following relation between their Young's moduli:

$$\frac{Y_{\text{Cur}}}{Y_{\text{pCe}}} \approx \frac{E_{r,\text{Cur}}}{E_{r,\text{pCe}}} \approx 2, \quad (2)$$

where the subscripts Cur and pCe designates $Cur^+$ and $pCe^+$ biofilms, respectively.
